# End-to-end differentiable learning of protein structure

**DOI:** 10.1101/265231

**Authors:** Mohammed AlQuraishi

## Abstract

Predicting protein structure from sequence is a central challenge of biochemistry. Co‐evolution methods show promise, but an explicit sequence‐to‐structure map remains elusive. Advances in deep learning that replace complex, human‐designed pipelines with differentiable models optimized end‐to‐end suggest the potential benefits of similarly reformulating structure prediction. Here we report the first end‐to‐end differentiable model of protein structure. The model couples local and global protein structure via geometric units that optimize global geometry without violating local covalent chemistry. We test our model using two challenging tasks: predicting novel folds without co‐evolutionary data and predicting known folds without structural templates. In the first task the model achieves state‐of‐the‐art accuracy and in the second it comes within 1‐2Å; competing methods using co‐evolution and experimental templates have been refined over many years and it is likely that the differentiable approach has substantial room for further improvement, with applications ranging from drug discovery to protein design.

## Main

Proteins are linear polymers that fold into very specific and ordered three dimensional conformations based on their amino acid sequence (Branden and Tooze, 1999; Dill, 1990). Understanding how this occurs is a foundational problem in biochemistry. Computational approaches to protein folding not only seek to make structure determination faster and less costly; they aim to understand the folding process itself. Existing computational methods fall into two broad categories (Gajda et al., 2011b, 2011a). The first category builds explicit sequence‐to‐structure maps using computational procedures to transform raw amino acid sequences into 3D structures. This includes physics‐based molecular dynamics simulations (Marx and Hutter, 2012), which are restricted by computational cost to small proteins, and fragment assembly methods (Gajda et al., 2011a), which find energy‐minimizing conformations by sampling statistically‐derived protein fragments. Fragment assembly usually achieves high accuracy only when homologous protein structures are used as templates. Such template‐based methods use one or more experimental structures—found through homology searches—as the basis for making predictions.

The second category of methods eschews explicit sequence‐to‐structure maps and instead identifies co‐evolving residues within protein families to derive residue‐residue contact maps, using co‐evolution as an indicator of contact in physical space (Hopf et al., 2014; Marks et al., 2011). With a large and diverse set of homologous sequences—typically tens to hundreds of thousands—co‐evolution methods can accurately predict contact maps (Juan et al., 2013). A correct contact map can guide fragment assembly methods to an accurate 3D structure 25‐50%of the time (Ovchinnikov et al., 2017). However, because co‐evolutionary methods do no construct a model of the relationship between individual sequences and structures, they are unable to predict structures for which no sequence homologs exist, as in new bacterial taxa or *de novo* protein design. Moreover, even for well‐characterized proteins, such methods are generally unable to predict the structural consequences of minor sequence changes such as mutations or indels, because they operate on protein families rather than individual sequences (they do however show promise in predicting the functional consequences of mutations (Hopf et al., 2017)). Thus, there remains a substantial need for new and potentially better approaches.

End‐to‐end differentiable deep learning has revolutionized computer vision and speech recognition (LeCun et al., 2015), but protein structure pipelines continue to resemble the ways in which computers tackled vision and speech prior to deep learning, by having many human‐engineered stages, each independently optimized (Xu and Zhang, 2012; Yang et al., 2015) (Fig. 1). End‐to‐end differentiable models replace all components of such pipelines with differentiable primitives to enable joint optimization from input to output. In contrast, use of deep learning for structure prediction has so far been restricted to individual components within a larger pipeline (Aydin et al., 2012; Gao et al., 2017; Li et al., 2017; Lyons et al., 2014), for example prediction of contact maps (Liu et al., 2017; Wang et al., 2016). This stems from the technical challenge of developing an end‐to‐end differentiable model that rebuilds the entire structure prediction pipeline using differentiable primitives. We have developed such a model by combining four ideas: (i) encoding protein sequence using a recurrent neural network, (ii) parameterizing (local) protein structure by torsional angles, to enable a model to reason over diverse conformations without violating their covalent chemistry, (iii) coupling local protein structure to its global representation via recurrent geometric units, and (iv) using a differentiable loss function to capture deviations between predicted and experimental structures. We find that the new approach outperforms other methods, including co‐evolution ones, when predicting novel folds even though it uses only primary sequences and position‐specific scoring matrices (PSSMs) that summarize individual residue propensities for mutation. We also find that when predicting known folds, the new approach is on average within 1‐2Å of other approaches, including template‐based ones, despite being template‐free.

**Fig. 1:**
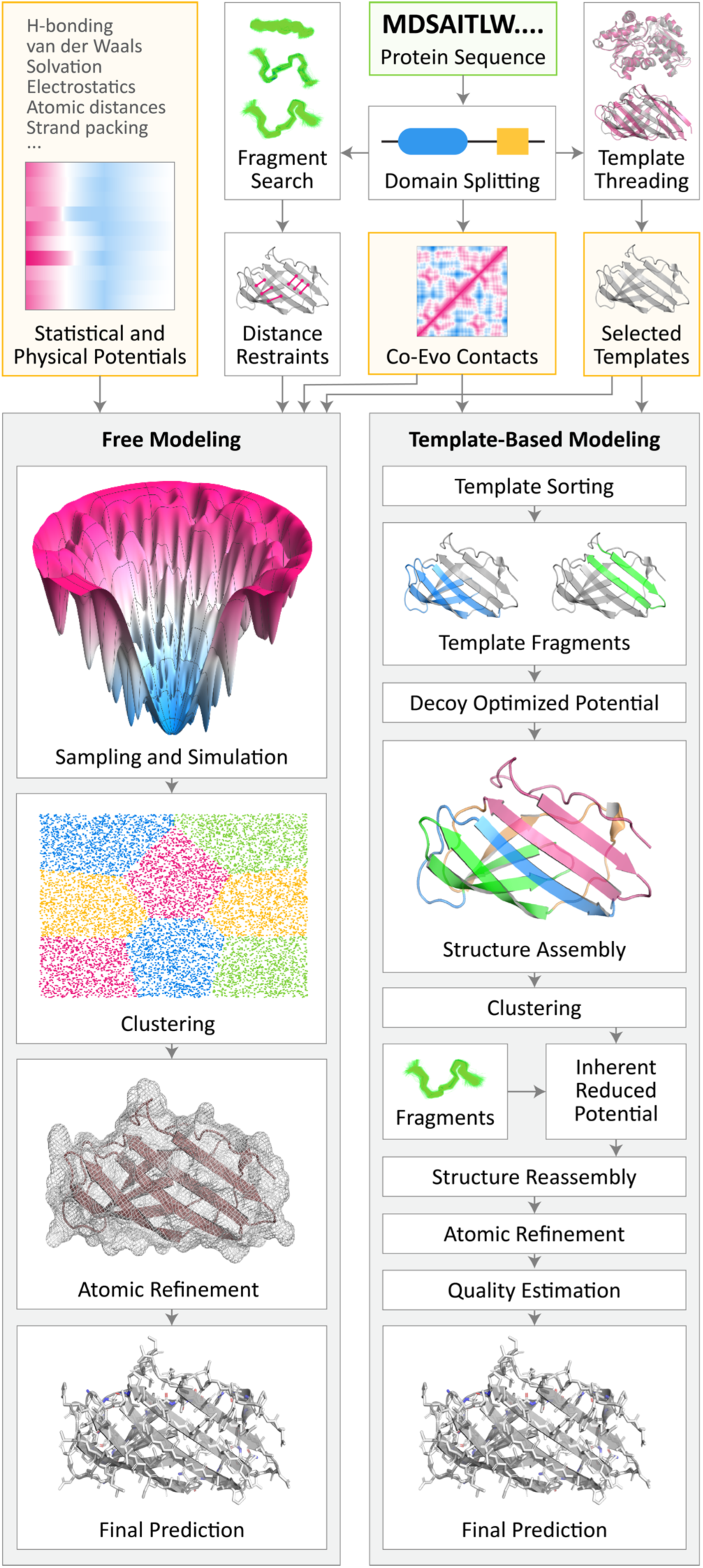
Conventional pipelines for protein structure prediction. Prediction process begins with query sequence (top, green box) whose constituent domains and co‐ evolutionary relationships are identified through multiple sequence alignments. In free modeling **(left)**, fragment libraries are searched to derive distance restraints which, along with restraints derived from co‐ evolutionary data, guide simulations that iteratively minimize energy through sampling. Coarse conformations are then refined to yield the final structure. In template‐based modeling **(right pipeline)**, the PDB is searched for templates. If found, fragments from one or more templates are combined to assemble a structure, which is then optimized and refined to yield the final structure. Orange boxes indicate sources of input information beyond query sequence, including prior physical knowledge. Diagram is modeled on the I‐Tasser and Quark pipelines (Zhang et al.).

### Recurrent geometric networks

Our model takes as input a sequence of amino acids and PSSMs and outputs a 3D structure. It is comprised of three stages—computation, geometry, and assessment—that we term a recurrent geometric network (RGN). The first stage is made of computational units that, for each residue position, integrate information about its amino acid and PSSM with information coming from adjacent units. By laying these units in a recurrent bidirectional topology (Fig. 2), the computations for each residue integrate information from residues upstream and downstream all the way to the N‐ and C‐terminus, covering the entire protein. By further stacking units in multiple layers (not shown), the model implicitly encodes a multi‐scale representation of proteins. Each unit outputs three numbers, corresponding to the torsional angles of the residue. We do not specify *a priori* how angles are computed. Instead, each unit’s computation is described by an equation whose parameters are optimized so that RGNs accurately predict structures.

**Fig. 2:**
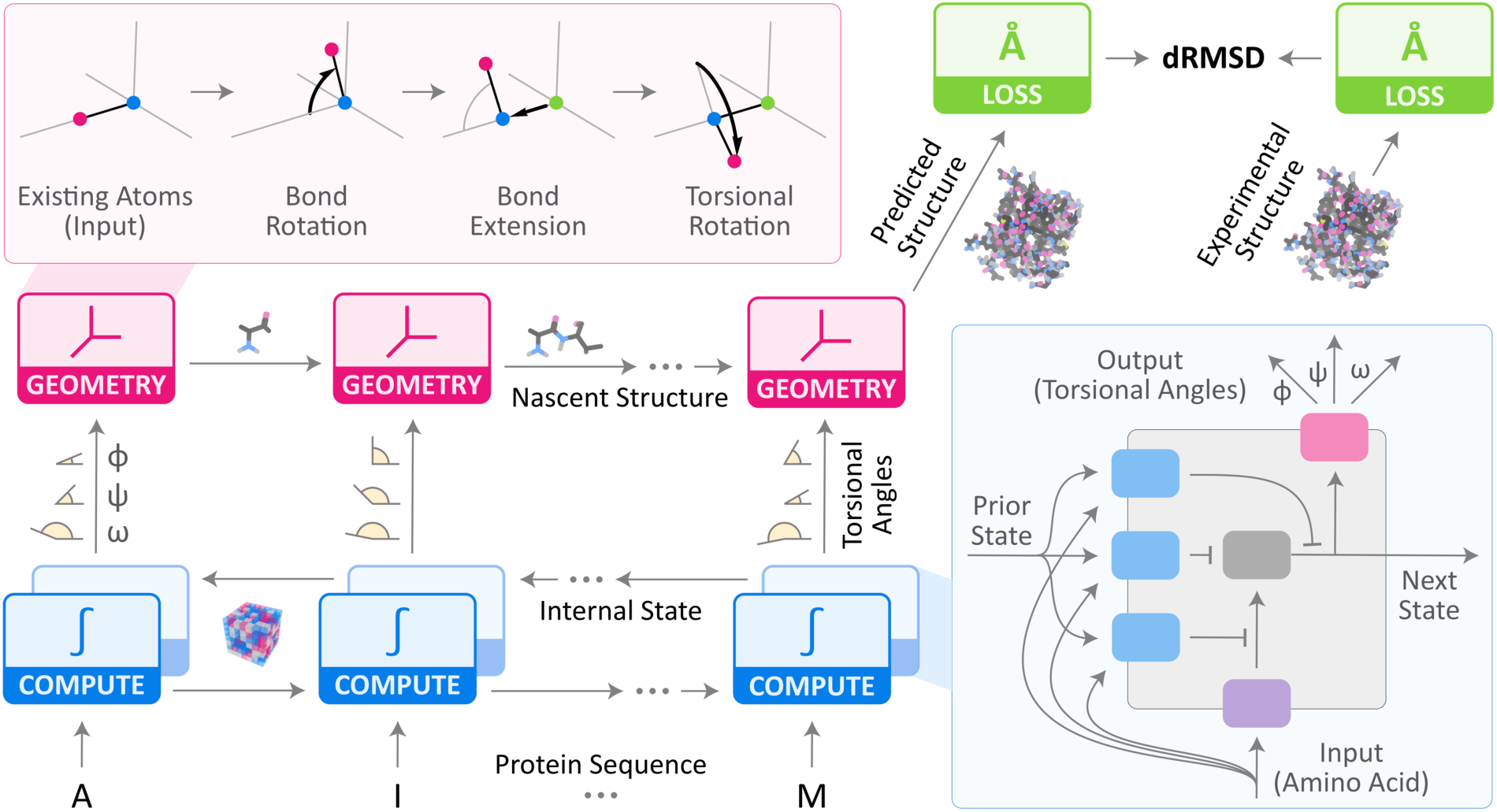
Recurrent geometric networks. Protein sequences are fed one residue at a time to the computational units of an RGN (bottom‐left), which compute an internal state that is integrated with the states of adjacent units. Based on these computations, torsional angles are predicted and fed to geometric units, which sequentially translate them into Cartesian coordinates to generate the predicted structure. dRMSD is used to measure deviation from experimental structures, serving as the signal for optimizing RGN parameters. **Top‐Left Inset:** Geometric units take new torsional angles and a partial backbone chain, and extend it by one residue. **Bottom‐Right Inset:** Computational units, based on Long Short‐Term Memory (LSTMs) (Hochreiter and Schmidhuber, 1997), use gating units (blue) to control information flow in and out of the internal state (gray), and angularization units (pink) to convert raw outputs into angles.

The second stage is made of geometric units that take as input the torsional angles for a given residue and the partially completed backbone resulting from the geometric unit upstream of it, and output a new backbone extended by one residue, which is fed into the adjacent downstream unit. The last unit outputs the completed 3D structure of the protein. During model training, a third stage computes deviations between predicted and experimental structures using the distance‐based root mean square deviation (dRMSD) metric. The dRMSD first computes pairwise distances between all atoms in the predicted structure and all atoms in the experimental one (separately), and then computes the root mean square of the distance between these sets of distances. Because dRMSD is distance‐based, it is invariant to reflections, which can lead RGNs to predict reflected structures (effectively wrong chirality) that must be corrected by a counter‐ reflection. RGN parameters are optimized to minimize the dRMSD between predicted and experimental structures using backpropagation (Goodfellow et al., 2016). Hyperparameters, which describe higher‐level aspects of the model such as the number of computational units, were determined through manual exploration of hyperparameter space. See Supplementary Text for a complete mathematical treatment.

### Assessment of model error

Machine learning models must be trained against as large a proportion of available data as possible to fit model parameters and then evaluated against a distinct test set to assess accuracy. Reliable evaluation is frequently complicated by unanticipated information leakage from the training set into the test set, especially for protein sequences which share an underlying evolutionary relationship. Partly to address this problem, the Critical Assessment of Protein Structure Prediction (CASP) (Moult et al., 1995) was organized to assess methods in a blinded fashion, by testing predictors using sequences of solved structures that have not been publicly released. To assess RGNs we therefore sought to recreate the conditions of past CASPs by assembling the ProteinNet datasets (Mohammed AlQuraishi, 2018). For every CASP from 7 through 12, we created a corresponding ProteinNet test set comprised of CASP structures, and a ProteinNet training set comprised of all sequences and structures publicly available prior to the start of that CASP. Using multiple CASP datasets enables a deeper and more thorough assessment that spans a broad range of dataset sizes than relying on the most recent CASP alone. We also adopted the CASP division of test structures into free modeling (FM) targets that assess prediction of novel folds, and template‐based (TBM and TBM‐hard) targets that assess prediction of folds with known homologs in the Protein Data Bank (PDB) (Bernstein et al., 1977). We set aside a subset of the training data as a validation set, to determine when to stop model training and to further insulate training and test data.

ProteinNet datasets were used for all analyses described here. RGN hyperparameters were fit by repeated evaluations on the ProteinNet 11 validation set followed by three evaluations on the ProteinNet 11 test set. Once chosen, the same hyperparameters were used to train models on ProteinNet 7‐12 training sets, with a single evaluation made at the end on each test set (excepting ProteinNet 11) to generate Table 1. Subsequently additional test set evaluations were made to generate Table S1, with one evaluation per number reported. No additional test set evaluations were made. Overall, this represents a rigorous approach to evaluation with the lowest possible risk of information leakage.

**Table 1:**
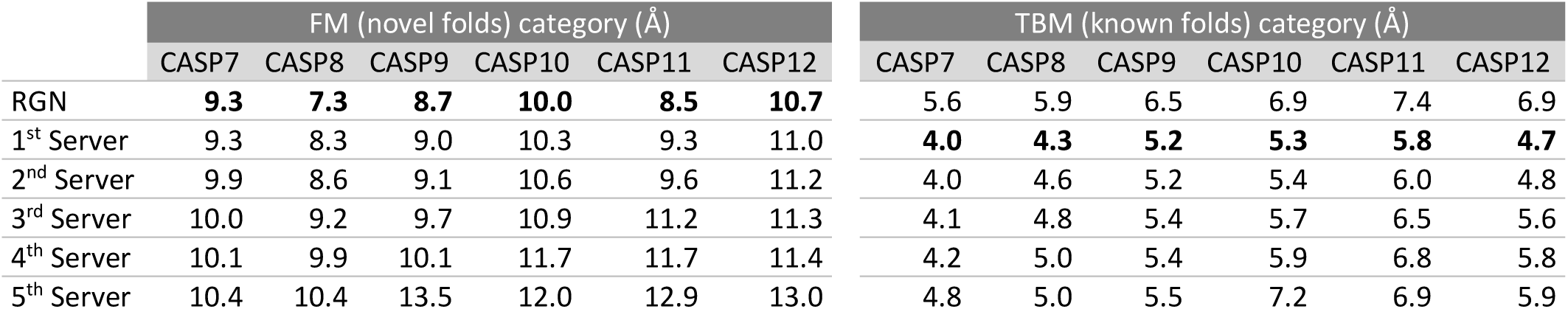
Comparative accuracy of RGNs using dRMSD. The average dRMSD (lower is better) achieved by RGNs and the top five servers at each CASP is shown for the novel folds **(left)** and known folds **(right)** categories. Numbers are based on common set of structures predicted by top 5 servers during each CASP. A different RGN was trained for each CASP, using the corresponding ProteinNet training set containing all sequences and structures available prior to the start of that CASP.

|  | FM (novel folds) category (Å) |  |  |  |  |  | TBM (known folds) category (Å) |  |  |  |  |  |
| --- | --- | --- | --- | --- | --- | --- | --- | --- | --- | --- | --- | --- |
|  | CASP7 | CASP8 | CASP9 | CASP10 | CASP11 | CASP12 | CASP7 | CASP8 | CASP9 | CASP10 | CASP11 | CASP12 |
| RGN | **9.3** | **7.3** | **8.7** | **10.0** | **8.5** | **10.7** | 5.6 | 5.9 | 6.5 | 6.9 | 7.4 | 6.9 |
| 1<sup>st</sup> Server | 9.3 | 8.3 | 9.0 | 10.3 | 9.3 | 11.0 | **4.0** | **4.3** | **5.2** | **5.3** | **5.8** | **4.7** |
| 2<sup>nd</sup> Server | 9.9 | 8.6 | 9.1 | 10.6 | 9.6 | 11.2 | 4.0 | 4.6 | 5.2 | 5.4 | 6.0 | 4.8 |
| 3<sup>rd</sup> Server | 10.0 | 9.2 | 9.7 | 10.9 | 11.2 | 11.3 | 4.1 | 4.8 | 5.4 | 5.7 | 6.5 | 5.6 |
| 4<sup>th</sup> Server | 10.1 | 9.9 | 10.1 | 11.7 | 11.7 | 11.4 | 4.2 | 5.0 | 5.4 | 5.9 | 6.8 | 5.8 |
| 5<sup>th</sup> Server | 10.4 | 10.4 | 13.5 | 12.0 | 12.9 | 13.0 | 4.8 | 5.0 | 5.5 | 7.2 | 6.9 | 5.9 |

### Predicting new folds without co‐evolution

We first assessed RGNs on a difficult task that has not consistently been achieved by any existing method: predicting novel protein folds without co‐evolutionary data. FM structures served as targets for this exercise. Table 1 compares the average dRMSD of RGN predictions on FM structures to the top five automated predictors in CASP 7‐12, known as “servers” in CASP parlance (“humans” are server/human‐expert pipelines—we do not compare against this group as our processing is automated). In Fig. 3a we break down the predictions by target against the top performing server and in Fig. 3c against the dRMSD distribution of all CASP servers.

**Fig. 3:**
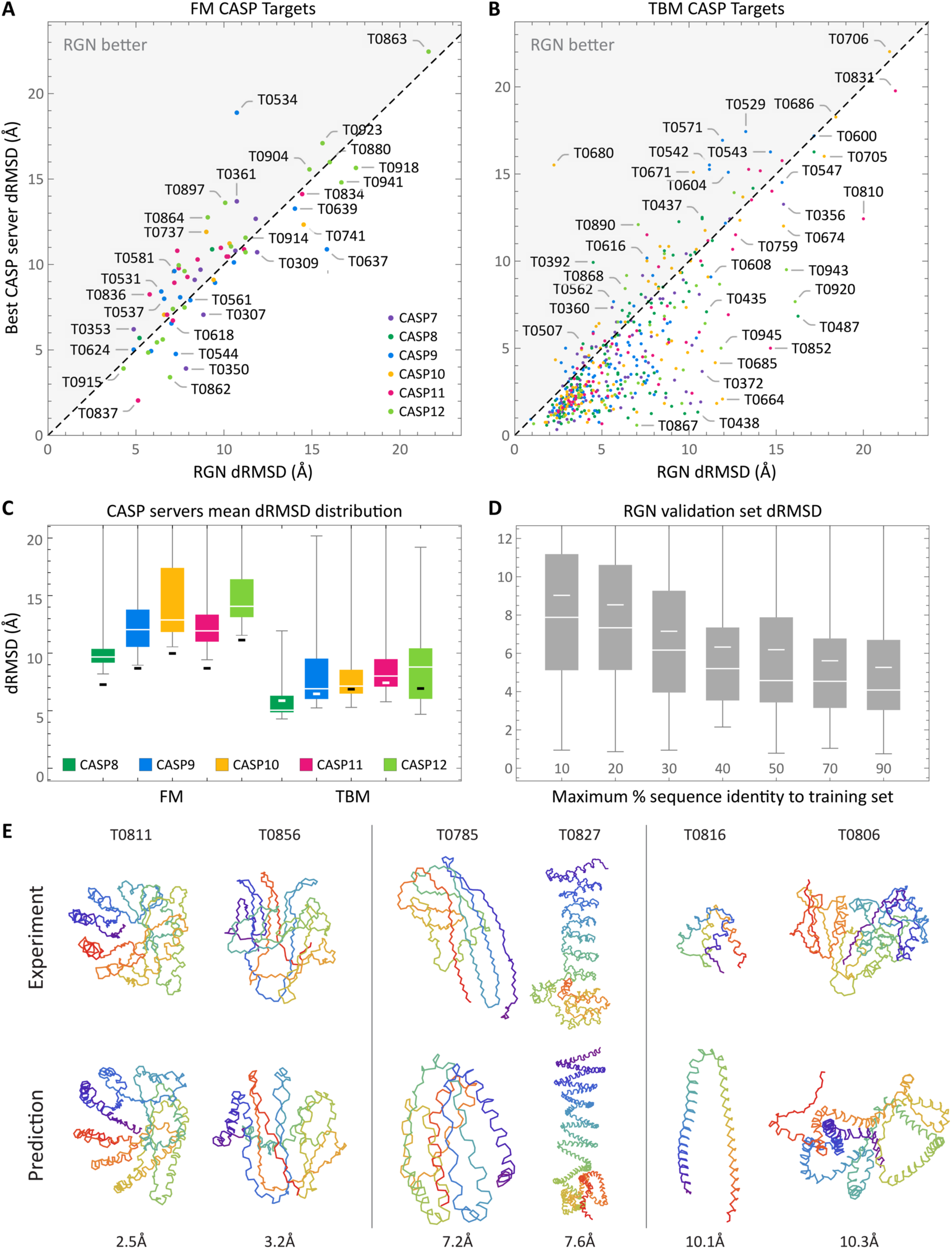
Results overview. Scatterplots of individual FM **(A)** and TBM **(B)** predictions made by RGN and top CASP server. Two TBM outliers (T0629 and T0719) were dropped for visualization purposes. **(C)** Distributions of mean dRMSD (lower is better, white is median) achieved by servers predicting all structures with >95%coverage at CASP 8‐12 are shown for FM (novel folds) and TBM (known folds) categories. Thick black (white on dark background) bars mark RGN dRMSD. CASP 7 is omitted due to lack of server metadata. **(D)** Distribution of RGN dRMSDs on ProteinNet validation sets grouped by maximum %sequence identity to training set over all CASPs (medians are wide white lines, means are short white lines.) **(E)** Traces of backbone atoms of well **(left)**, fairly **(middle)**, and poorly **(right)** predicted RGN structures are shown **(bottom)** along with their experimental counterparts **(top)**. CASP identifier is displayed above each structure and dRMSD below. A color spectrum spans each protein chain to aid visualization.

On all CASPs, RGNs had the best performance, even compared to servers that use co‐evolution data (in CASP 11 (Kryshtafovych et al., 2016; Ovchinnikov et al., 2016) and CASP 12 (Schaarschmidt et al., 2017)). RGNs outperformed other methods at both short and long, multi‐domain proteins, suggesting their performance is not limited to one regime (e.g. short single domain proteins), despite having no explicit knowledge of domain boundaries. While the margin between RGNs and the next best server is small for most CASPs, such small gaps are representative of the differences between the top five performers in Table 1. In general, small gains in accuracy at the top end are difficult, with only minimal gains obtained over a ten‐year time span from CASP 6 to CASP 11 (Kryshtafovych et al.). More substantial gains were seen in CASP 12 due to the use of co‐evolutionary information (Moult John et al., 2018), but RGNs match these advances without using co‐evolutionary data and by operating in a fundamentally distinct and complementary way. The accuracy gap between RGNs and other servers is highest on CASP 11, which benefits from having the RGN hyperparameters fit on the ProteinNet11 validation set, suggesting similar gains may be had by optimizing RGN hyperparameters for each dataset (this would not correspond to overfitting, as only the validation set is used to fit hyperparameters, but would require substantially more compute resources for training.) ProteinNet datasets of earlier CASPs are smaller which may have also reduced accuracy. To assess the contribution of dataset size to model error, we used RGNs trained on earlier ProteinNet datasets to predict later CASP test sets (Table S1). As expected, accuracy drops as datasets shrink.

The dRMSD metric does not require structures to be pre‐aligned, and is consequently able to detect regions of high local concordance even when global concordance is poor. Because dRMSD assesses predictions at all length scales however, it penalizes large global deviations in proportion to their distance, which can result in very high error for far apart regions. To obtain a complementary assessment of model accuracy, we also tested RGNs using TM scores (Zhang and Skolnick, 2004), which are defined by the following equation:

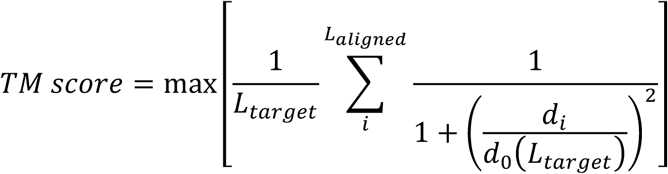

where *L*_*target*_ and *L*_*aligned*_ are the lengths of the full protein and the aligned region, respectively, *d*_*i*_ is the distance between the i^th^residues in the experimental and predicted structures, and 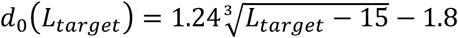 is used to normalize scores. TM scores do require structures to be pre‐aligned, and thus can penalize predictions with high local concordance if a global alignment cannot be found, but they are less sensitive to large deviations because they only compute error over the aligned regions. TM scores range from 0 to 1, with a score of < 0.17 corresponding to a random unrelated protein, and > 0.5 generally corresponding to the same protein fold (Xu and Zhang, 2010). Since TM scores are not invariant to reflections, we compute them for both the original and reflected RGN structures and use the higher of the two. Table S2 compares TM scores of RGN predictions to CASP servers. In general, RGNs rank among the top five servers, but do not consistently outperform all other methods as they do on dRMSD, possibly reflecting the lack of partial credit assignment by TM scores.

### Predicting known folds without templates

We next assess RGNs on predicting known protein folds without experimental templates, a challenging task that provides an advantage to template‐based methods (Zhou et al., 2010). TBM structures served as targets for this purpose. Table 1 and Table S2 compare RGN predictions to top CASP servers using dRMSD and TM score, respectively, while Fig. 3b breaks down predictions by target and Fig. 3c shows the distribution over all CASP servers. A representative sampling of the full quality spectrum of FM and TBM predictions is shown in Fig. 3e. In general, RGNs underperform the very top CASP servers, all of which use templates, although ∼60%of predictions are within 1.5Å of the best‐performing server.

Since RGNs do not use templates, this suggests that they learn generalizable aspects of protein structure, and their improved accuracy on TBM targets relative to FM reflects denser sampling in TBM regions of protein space. To investigate this possibility, we partitioned ProteinNet validation sets into groups based on maximum sequence identity to the training set, and computed dRMSDs within each group across CASPs 7‐12 (Fig. 3d) and by individual CASP (Fig. S1). RGN performance robustly transfers to sequences with >40% sequence identity, predicting structures with a median dRMSD of ∼5Å, and then begins to deteriorate. There was little difference in dRMSD between 50% and 90% sequence identity, with substantial error remaining at 90%, which is suggestive of underfitting.

Template‐based methods are particularly accurate where template and query sequences overlap, and are inaccurate where they do not; unfortunately, non‐overlapping regions are often the regions of high biological interest. Errors in these critical non‐overlapping regions can be masked by large overlapping regions, inflating overall accuracy (Contreras‐Moreira et al., 2005; Dill and MacCallum, 2012; Liu et al.; Perez et al., 2016). To determine whether RGNs suffer from similar limitations, we split TBM domains into short fragments ranging in size from 5 to 50 residues and computed the RMSD for every fragment (with respect to the experimental structure) from the best template, the best CASP prediction, and the RGN prediction (Fig. 4). We found CASP predictions to be correlated (average *R*^2^= 0.44) with template quality across length scales as previously reported (Kryshtafovych et al.), while RGN predictions were not (average *R*^2^= 0.06). Thus RGNs perform equally well on regions of proteins with experimental templates and on those without.

**Fig. 4:**
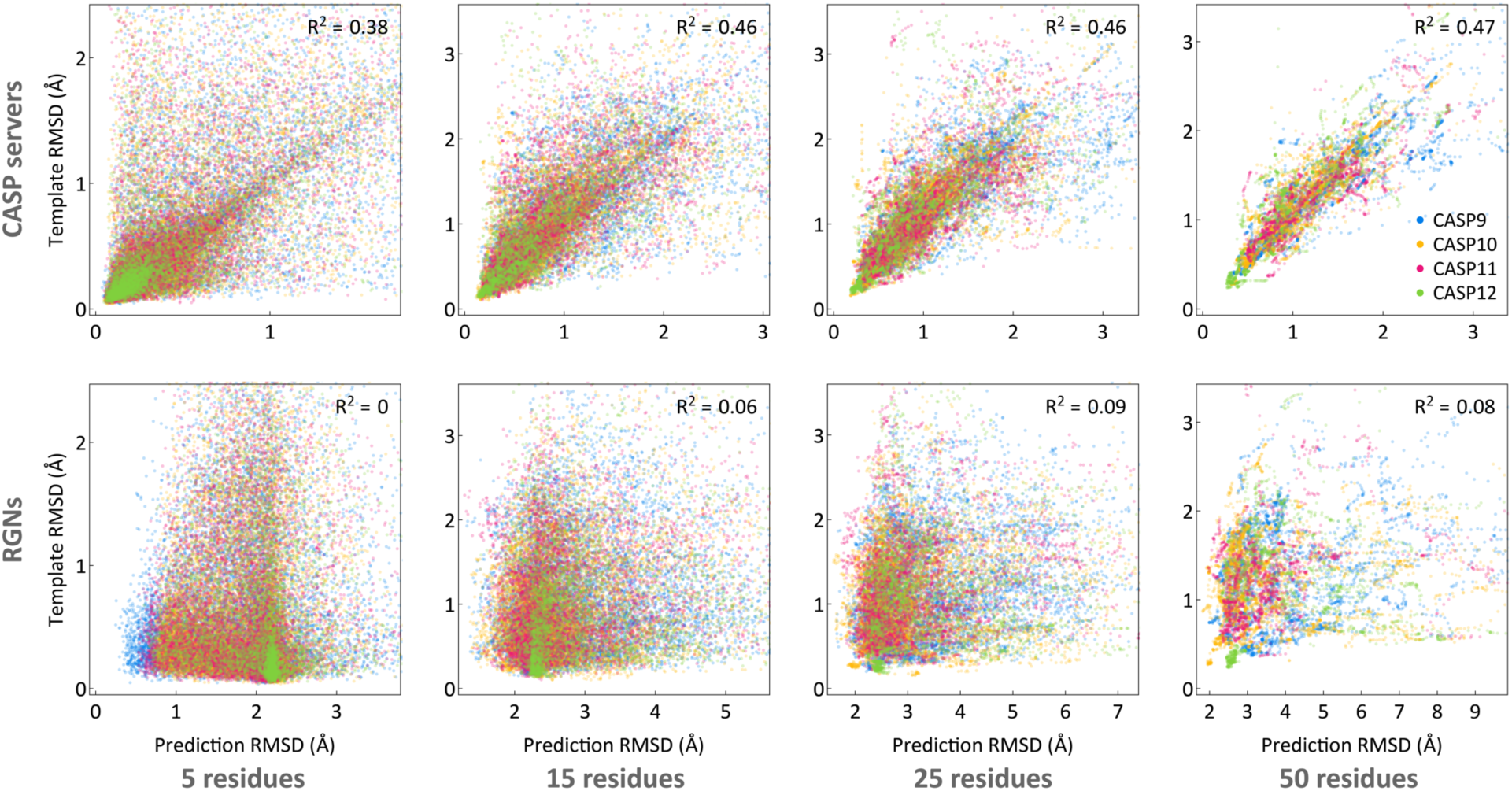
Correlation between prediction accuracy and template quality. Scatterplots of fragment RMSDs, ranging in size from 5 to 50 residues, comparing the best CASP templates to the best CASP server predictions **(top)** and RGN predictions **(bottom)**. TBM domains were used (excluding TBM‐hard which do not have good templates), and only templates and predictions covering >85%of full domain sequences were considered. Templates and predictions were selected based on global dRMSD with respect to experimental structure. CASP 7 and 8 are omitted due to lack of full template information.

### RGNs learn an implicit representation of protein fold space

Applications of deep learning in sensory domains often result in models whose internal representation of the data is interpretable, e.g. placing semantically similar words nearby in a natural language model. To ascertain whether RGNs behave similarly, we extracted the internal state of their computational units after processing each protein sequence in the ProteinNet12 training set. For each protein, we obtained multiple high‐dimensional vectors, one per layer / direction of the RGN. We then used linear dimensionality reduction techniques to visualize these vectors in two dimensions, separately for each layer / direction (Fig. 5a), and by concatenating all layers together (Fig. 5b). When we color each protein (dot) according to the fraction of secondary structure present in its original PDB structure, clear visual patterns emerge (Fig. 5b). This is notable because secondary structure was neither used as input to aid model prediction nor as an output signal to guide training; i.e. the model was not explicitly encoded with the concept of secondary structure, yet it uses secondary structure as the dominant factor in shaping its representation of protein fold space.

**Fig. 5:**
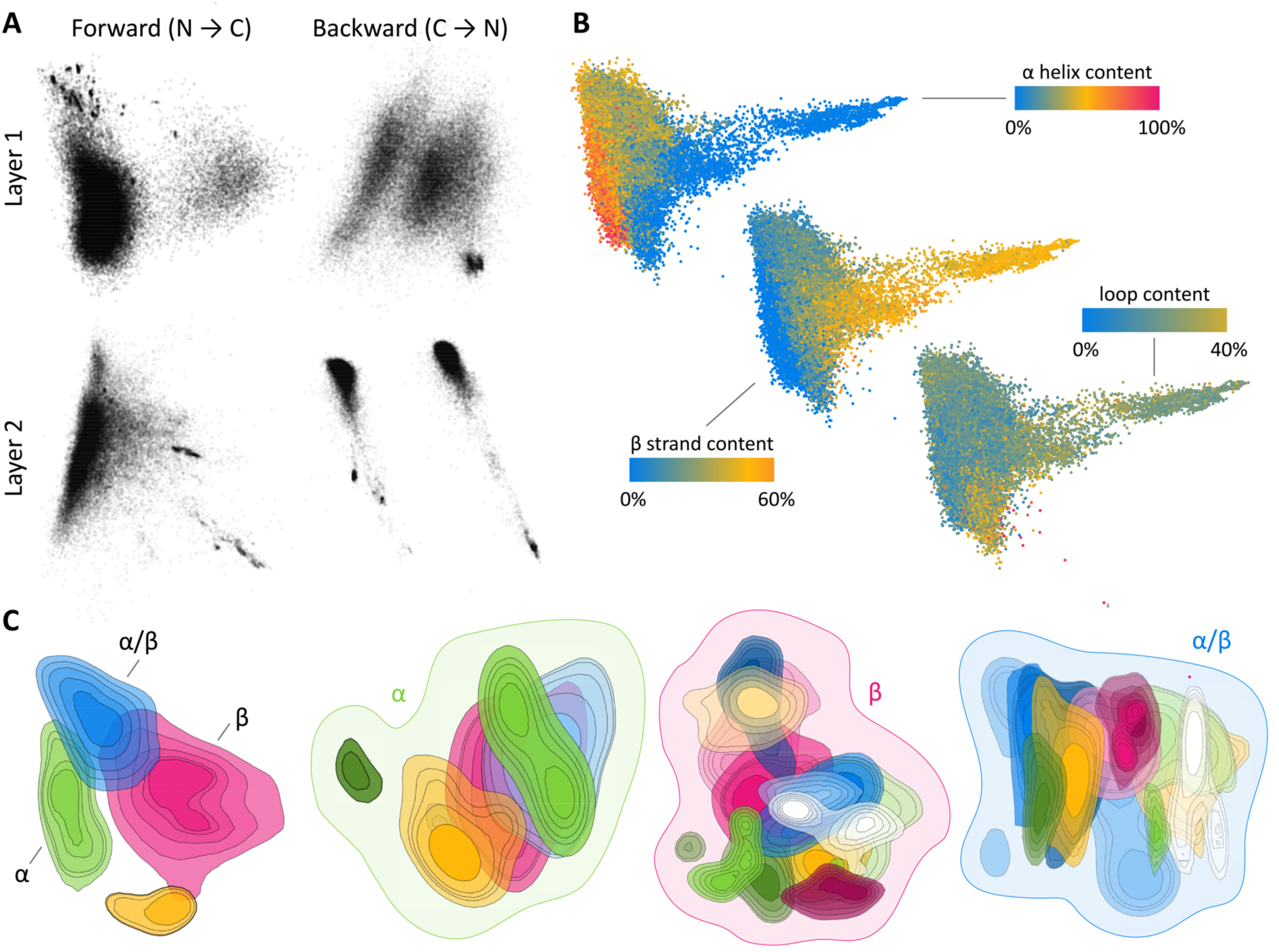
The latent space of RGNs. 2D projection of the separate **(A)** and combined **(B)** internal state of all RGN computational layers, with dots corresponding to individual protein sequences in the ProteinNet12 training set. **(B)** Proteins are colored by fractional secondary structure content, as determined by annotations of original protein structures. **(C)** Contour plots of the probability density (50‐90%quantiles) of proteins belonging to categories in the topmost level of the CATH hierarchy **(first from left)** and proteins belonging to categories in the second‐level CATH classes of “Mainly Alpha” **(second)**, “Mainly Beta” **(third)**, and “Alpha Beta” **(fourth)**. Distinct colors correspond to distinct CATH categorizations; see Fig. S2‐S5 for complete legends. The topmost CATH class “Few Secondary Structures” is omitted because it has no subcategories.

We next used the CATH database (Dawson et al., 2017), which hierarchically classifies proteins into structural families, to partition data points into CATH classes and visualize their distribution in RGN space. At the topmost CATH level, divided into “Mainly Alpha”, “Mainly Beta”, “Alpha Beta”, and “Few Secondary Structures”, we see clearly demarcated regions for each class (represented by differently colored contour plots), with “Alpha Beta” acting unsurprisingly as the bridge (leftmost panel in Fig. 5c.) We then reapplied dimensionality reduction to data in each class and visualized the distributions of their respective second‐level CATH categories (three right panels in Fig. 5c.) We again see contiguous regions for each category, albeit with greater overlap, likely owing to the continuous nature of protein structure space and reduction of RGN space to just two dimensions. These visualizations suggest RGNs are learning a useful representation of protein sequence space that may yield insights into the nature of protein structure space.

### RGNs are 6‐7 orders of magnitude faster than existing methods

Existing structure prediction pipelines are multi‐staged (Fig. 1), first detecting domains that can be separately modelled, and running multiple algorithms to estimate secondary structure propensities, solvent accessibility, and disordered regions. Co‐evolutionary methods use multiple sequence alignments to predict contact maps, and template‐based methods search the PDB for templates. Their predictions are converted into geometric constraints to guide a conformation sampling process, where fragments are swapped in and out of putative structures to minimize an expertly‐derived energy model. Due to this complexity, prediction times range from several hours to days, and require codebases as large as several million lines of code (Leaver‐Fay et al., 2011).

In contrast, a trained RGN model is a single mathematical function that is evaluated once per prediction. Computation of this function implicitly carries out domain splitting, property finding, energy minimization, and conformational sampling simultaneously. We found that 512 concurrent RGN‐based predictions, with sequence length ∼700, can be made in ∼5.4 seconds on a single GPU, i.e. ∼10 milliseconds / structure. Table 2 compares training and prediction speeds of RGNs to established methods that rely heavily on simulation with limited learning (first row), and co‐evolution‐based contact prediction methods that rely on learning (second row), combined with CONFOLD (Adhikari et al., 2015) to convert predicted contact maps into tertiary structures. While training RGNs can take weeks to months, once trained, they make predictions 6‐7 orders of magnitude faster than existing pipelines. This speed enables new types of applications, such as the integration of structure prediction within docking and virtual screening in which ligand‐aware RGNs could output distinct protein conformations in response to distinct ligand poses.

**Table 2:**
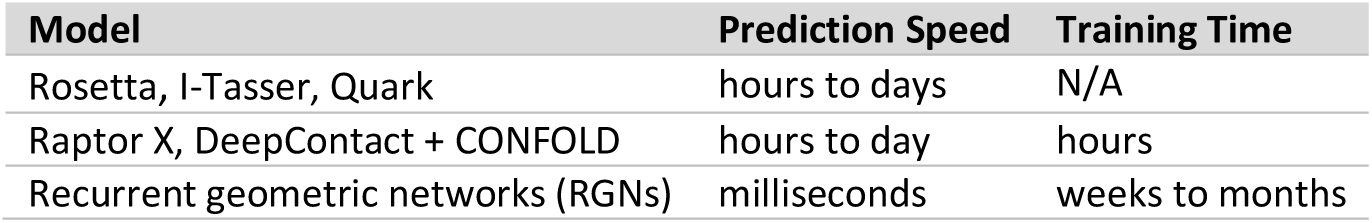
Prediction and training speeds of structure prediction methods. Top row corresponds to the most complex and established set of methods, which rely heavily on simulation and sampling, and typically have only a minimal learning component. Second row corresponds to co‐evolution‐based contact prediction methods, which rely on a learning procedure, plus the CONFOLD method to convert predicted contact maps into tertiary structures.

## Discussion

A key limitation of explicit sequence‐to‐structure maps, including molecular dynamics and fragment assembly, is a reliance on fixed energy models that do not learn from data; a second limitation is the exclusive use of single‐scale atomic or residue‐level representations. In contrast, modern co‐evolution methods leverage learning and multi‐scale representations to substantially improve performance (Liu et al., 2017; Wang et al., 2016). RGNs go one step further by building a fully differentiable map extending from sequence to structure with all of the steps in existing prediction pipelines implicitly encoded and learnable from data. Through their recurrent architecture, RGNs can capture sequence‐structure motifs and multiple scales from residues to domains (Alva et al., 2015; Ponting and Russell, 2002). When tracking structure prediction during RGN training (Movie S1), RGNs appear to first learn global aspects of protein folds, then refine their predictions to generate more accurate local structure.

RGNs are multi‐representational, operating on three distinct parameterizations of protein structure. The first is torsional, capturing local relationships between atoms with bond lengths and angles held fixed, and torsional angles as the immediate outputs of computational units. This virtually guarantees that predictions are structurally correct at a local level. The second is Cartesian, built by geometric units and capturing the global coordination of multiple atoms in 3D space, the catalytic triad of an enzyme’s active site for example, even if the residues are distant along the protein chain. Future augmentations—e.g. 3D convolutional networks that operate directly on the Cartesian representation—may further improve the detection and quality of long‐range interactions. The third parameterization, built in the dRMSD stage, is the matrix of inter‐ atomic distances, and is simultaneously local and global. It is useful for optimizing RGN parameters *de novo*, as we have used it, but can also be used to incorporate prior knowledge expressible in terms of atomic distances; such knowledge includes physical features (e.g. electrostatics) and statistical data on interactions (e.g. evolutionary couplings).

One limitation of current RGNs is their reliance on PSSMs, which we have found to be helpful to achieving high accuracy predictions. PSSMs are much weaker than multiple sequence alignments as they are based on single residue mutation frequencies and ignore how each residue mutates in response to all other residues. Co‐evolutionary couplings require pairwise frequencies, resulting in quadratically rather than linearly scaling statistical cost. Nonetheless, removing PSSMs and relying exclusively on raw sequences could robustify RGNs for many applications, including prediction of genetic variants. Achieving this may require more data‐efficient model architectures. For protein design, RGNs can be used as is, by fixing the desired structure and optimizing the raw sequence and PSSMs to match it (i.e. by computing derivatives of the inputs—as opposed to model parameters—with respect to the dRMSD between predicted and desired structures.) Co‐evolution methods do not have this capability as their inputs are the inter‐residue couplings themselves, making the approach circular.

The history of protein structure prediction suggests that new methods complementary to existing ones are eventually incorporated into hybrids. RGNs have this benefit, being an almost entirely complementary modeling approach. For example, structural templates or co‐evolutionary information could be incorporated as priors in the distance‐based parameterization or even as raw inputs for learning. RGNs can also include secondary structure predicted by other algorithms. This is likely to be advantageous since the RGNs described here often predict global fold correctly but do less well with secondary structure (e.g. T0827 in Fig. 3e). RGNs can also be made to predict side‐chain conformations, by outputting a branched curve in lieu of the current linear curve, and are applicable to a wide range of other polymers (e.g. RNA tertiary structure.) Our demonstration that state of the art performance in structure prediction can be achieved using an end‐to‐end differentiable model will make available to protein folding and biophysics very rapid improvements in machine learning across a wide range of scientific and technical fields. We predict that hybrid systems using deep learning, co‐evolution as priors, and physics‐based approaches for refinement will soon solve the long‐standing problem of accurate and efficient structure prediction. It is also possible that the use of neural network probing techniques (Alain and Bengio, 2016; Koh and Liang, 2017; Nguyen et al., 2016; Shrikumar et al., 2017; Simonyan et al., 2013) with RGNs will provide new insight into the physical chemistry of folding and the sorts of intermediate structures that proteins use to sample conformational space.

## Acknowledgments

We are indebted to Peter Sorger for his mentorship and support, and thank him for extensive editorial feedback on this manuscript. We thank Jasper Snoek and Adrian Jinich for their editorial comments and many helpful discussions, and Uraib Aboudi, Ramy Arnaout, Karen Sachs, Michael Levitt, and Nazim Bouatta for their feedback. We also thank Martin Steinegger and Milot Mirdita for their help with using the HHblits and MMseqs2 packages, Sergey Ovchinnikov for discussions about the manuscript and help with metagenomics sequences, Andriy Kryshtafovych for his help with CASP structures, Sean Eddy for his help with using the JackHMMer package, and Raffaele Potami, Amir Karger, and Kristina Holton for their help with using the HPC resources at Harvard Medical School. We gratefully acknowledge the support of NVIDIA Corporation with the donation of the Titan Xp GPUs used for this research.

## Supplementary Text

### Model

We featurize a protein of length *L* as a sequence of vectors (*x*_1_, …,*x*_*L*_) where *x*_*t*_ ∈ ℝ^*d*^ for all *t*. The dimensionality *d* is 41, where 20 dimensions are used as a one‐hot indicator of the amino acid residue at a given position, another 20 dimensions are used for the PSSM of that position, and 1 dimension is used to encode the information content of the position. The PSSM values are sigmoid transformed to lie between 0 and 1. The sequence of input vectors are fed to an LSTM (Hochreiter and Schmidhuber, 1997), whose basic formulation is described by the following set of equations.

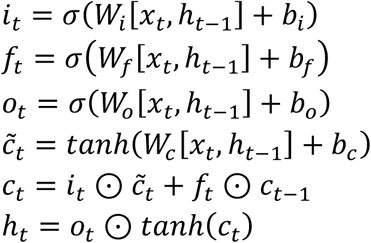

*W*_*i*_, *W*_*f*_, *W*_*o*_, *W*_*c*_, are weight matrices, *b*_*i*_, *b*_*f*_, *b*_*o*_, *b*_*c*_, are bias vectors, *h*_*t*_ and *c*_*t*_ are the hidden and memory cell state for residue *t*, respectively, and ⊙ is element‐wise multiplication. We use two LSTMs, running independently in opposite directions (1 to *L* and *L* to 1), to output two hidden states 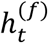 and 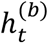 for each residue position *t* corresponding to the forward and backward directions. Depending on the RGN architecture, these two hidden states are either the final outputs states or they are fed as inputs into one or more LSTM layers.

The outputs from the last LSTM layer form a sequence of a concatenated hidden state vectors 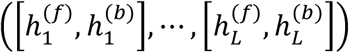. Each concatenated vector is then fed into an angularization layer described by the following set of equations:

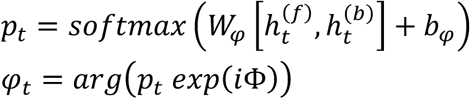

*W*_*φ*_ is a weight matrix, *b*_*φ*_ is a bias vector, Φ is a learned alphabet matrix, and *arg* is the complex‐valued argument function. Exponentiation of the complex‐valued matrix *i*Φ is performed element‐wise. The Φ matrix defines an alphabet of size *m* whose letters correspond to triplets of torsional angles defined over the 3‐torus. The angularization layer interprets the LSTM hidden state outputs as weights over the alphabet, using them to compute a weighted average of the letters of the alphabet (independently for each torsional angle) to generate the final set of torsional angles *φ*_*t*_ ∈ *S*^1^ × *S*^1^ × *S*^1^ for residue *t* (we are overloading the standard notation for protein backbone torsional angles, with *φ*_*t*_ corresponding to the (*ψ*, *φ*, *ω*) triplet). Note that *φ*_*t*_ may be alternatively computed using the following equation, where the trigonometric operations are performed element‐wise:

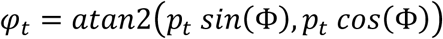

In general, the geometry of a protein backbone can be represented by three torsional angles φ, ψ, and ω that define the angles between successive planes spanned by the N, C^α^, and C’ protein backbone atoms (Ramachandran et al., 1963). While bond lengths and angles vary as well, their variation is sufficiently limited that they can be assumed fixed. Similar claims hold for side chains as well, although we restrict our attention to backbone structure. The resulting sequence of torsional angles (*φ*_1_, …, *φ*_*L*_) from the angularization layer is fed sequentially, along with the coordinates of the last three atoms of the nascent protein chain (*c*_1_,…*c*_3t_), into recurrent geometric units that convert this sequence into 3D Cartesian coordinates, with three coordinates resulting from each residue, corresponding to the N, C^α^, and C’ backbone atoms. Multiple mathematically‐equivalent formulations exist for this transformation; we adopt one based on the Natural Extension Reference Frame (Strauss et al., 2005), described by the following set of equations:

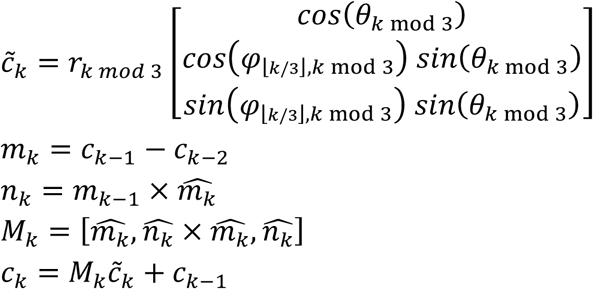

Where *r*_*k*_ is the length of the bond connecting atoms *k* - 1 and *k, θ*_*k*_ is the bond angle formed by atoms *k* - 2, *k* - 1 and *k, φ* _⌊*k*/3⌋,*k* mod 3_ is the predicted torsional angle formed by atoms *k* - 2 and *k* - 1, *c*_*k*_ is the position of the newly predicted atom *k*, 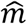 is the unit‐normalized version of *m*, and × is the cross product. Note that *k* indexes atoms 1 through 3*L*, since there are three backbone atoms per residue. For each residue *t* we compute *c*_3*t*–2,_ *c*_3*t*–1_, and *c*_3*t*_ using the three predicted torsional angles of residue *t*, specifically 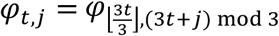 for *j* = {0,1,2}. The bond lengths and angles are fixed, with three bond lengths (*r*_0_, *c*_1_, *c*_2_) corresponding to N‐C^α^, C^α^‐C’, and C’‐N, and three bond angles (*θ*_0_, *θ*_1_, *θ*_2_) corresponding to N‐C^α^‐C’, C^α^‐C’‐N, and C’‐N‐C^α^. As there are only three unique values we have *r*_*k*_ = *r*_*k* mod 3_ and *θ*_*k*_ = *θ*_*k* mod 3_. In practice we employ a modified version of the above equations which enable much higher computational efficiency, described in an upcoming paper.

The resulting sequence (*c*_1_,…,*c*_3*L*_) fully describes the protein backbone chain structure and is the model’s final predicted output. For training purposes a loss is necessary to optimize model parameters. We use the *dRMSD* metric as it is differentiable and captures both local and global aspects of protein structure. It is defined by the following set of equations:

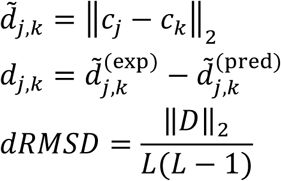

Where {*d*_*j,k*_} are the elements of matrix *D*, and 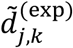 and 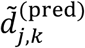 are computed using the coordinates of the experimental and predicted structures, respectively. In effect, the *dRMSD* computes the *l*_2_ ‐norm of the distances over distances, by first computing the pairwise distances between all atoms in both the predicted and experimental structures individually, and then computing the distances between those distances. For most experimental structures, the coordinates of some atoms are missing. They are excluded from the dRMSD by not computing the differences between their distances and the predicted ones.

<u>Hyperparameters</u>

RGN hyperparameters were manually fit, through sequential exploration of hyperparameter space, using repeated evaluations on the ProteinNet11 validation set and three evaluations on ProteinNet11 test set. Once chosen the same hyperparameters were used to train RGNs on ProteinNet7‐12 training sets. The validation sets were used to determine early stopping criteria, followed by single evaluations on the ProteinNet7‐12 test sets to generate the final reported numbers (excepting ProteinNet11).

The final model consisted of two bidirectional LSTM layers, each comprised of 800 units per direction, and in which outputs from the two directions are first concatenated before being fed to the second layer. Input dropout set at 0.5 was used for both layers, and the alphabet size was set to 60 for the angularization layer. Inputs were duplicated and concatenated; this had a separate effect from decreasing dropout probability. LSTMs were random initialized with a uniform distribution with support [–0.01, 0.01] while the alphabet was similarly initialized with support [−*π*, *π*]. ADAM was used as the optimizer, with a learning rate of 0.001, *β*_1_ = 0.95 and *β*_2_ = 0.99 and a batch size of 32. Gradients were clipped using norm rescaling with a threshold of 5.0. The loss function used for optimization was length‐normalized dRMSD (i.e. dRMSD divided by protein length), which is distinct from the standard dRMSD we use for reporting accuracies.

RGNs are very seed sensitive. As a result, we used a milestone scheme to restart underperforming models early. If a dRMSD loss milestone is not achieved by a given iteration, training is restarted with a new initialization seed. Table S3 summarizes the milestones, which were determined based on preliminary runs. In general, 8 models were started and, after surviving all milestones, were run for 250k iterations, at which point the lower performing half were discarded, and similarly at 500k iterations, ending with 2 models that were usually run for ∼2.5M iterations. Once validation error stabilized we reduced the learning rate by a factor of 10 to 0.0001, and run for a few thousand additional iterations to gain a small but detectable increase in accuracy before ending model training.

## Dataset

We use the ProteinNet dataset for all analyses performed, which is described in detail elsewhere (Mohammed AlQuraishi, 2018). ProteinNet recreates the conditions of past CASP assessments by restricting the set of sequences (for building PSSMs) and structures used to those available prior to the start of each CASP assessment. Each ProteinNet entry is comprised of two inputs, the raw protein sequence, represented by a one‐hot vector, and the protein’s PSSM and information content profiles, derived using 5 iterations of JackHMMer with an e‐value threshold of 10^‐10^. PSSM values are normalized to lie between 0 and 1. The output for each ProteinNet entry is comprised of the Cartesian coordinates of the protein’s backbone atoms, annotated by metadata denoting which atoms are missing from the experimental structure. These atoms are excluded from the dRMSD loss calculation, which enables use of partially resolved experimental structures that would otherwise be excluded from the dataset.

For ProteinNet7‐11, the publicly available CASP structures were used as test sets. For ProteinNet12, the publicly available CASP12 structures are incomplete, as some structures are still embargoed. We obtained a private set of structures from the CASP organizers that includes all structures used in CASP12 (except one or two), and we used this set for model assessment. For training all RGN models, the 90%“thinning” version of ProteinNet was used.

---

**Fig. S1.**
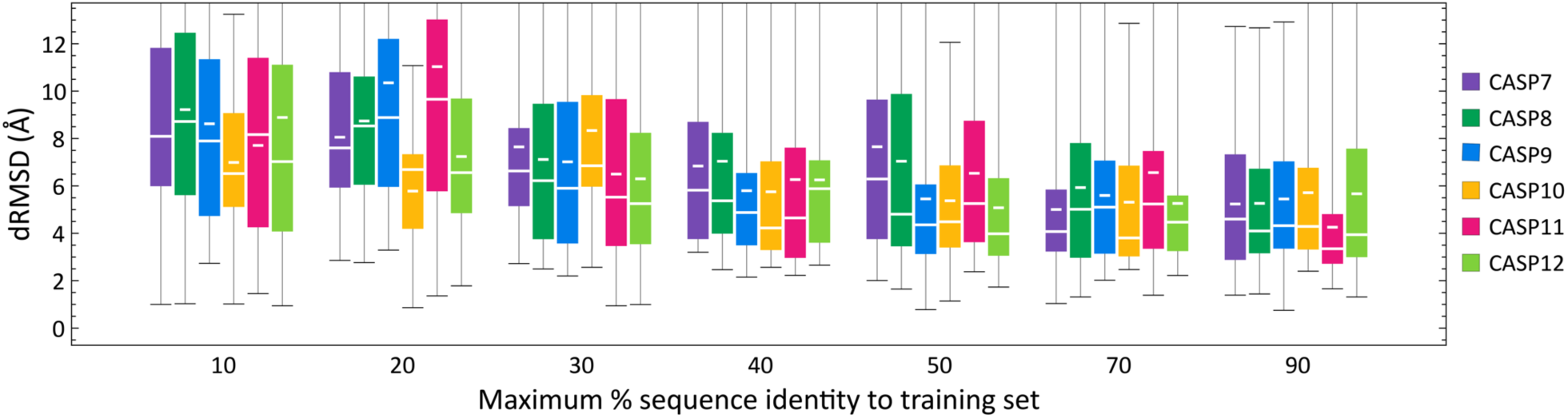
Distribution of dRMSDs of ProteinNet validation sets grouped by maximum %sequence identity to training set and broken down by each CASP (medians are wide white lines, means are short white lines).

**Fig. S2.**
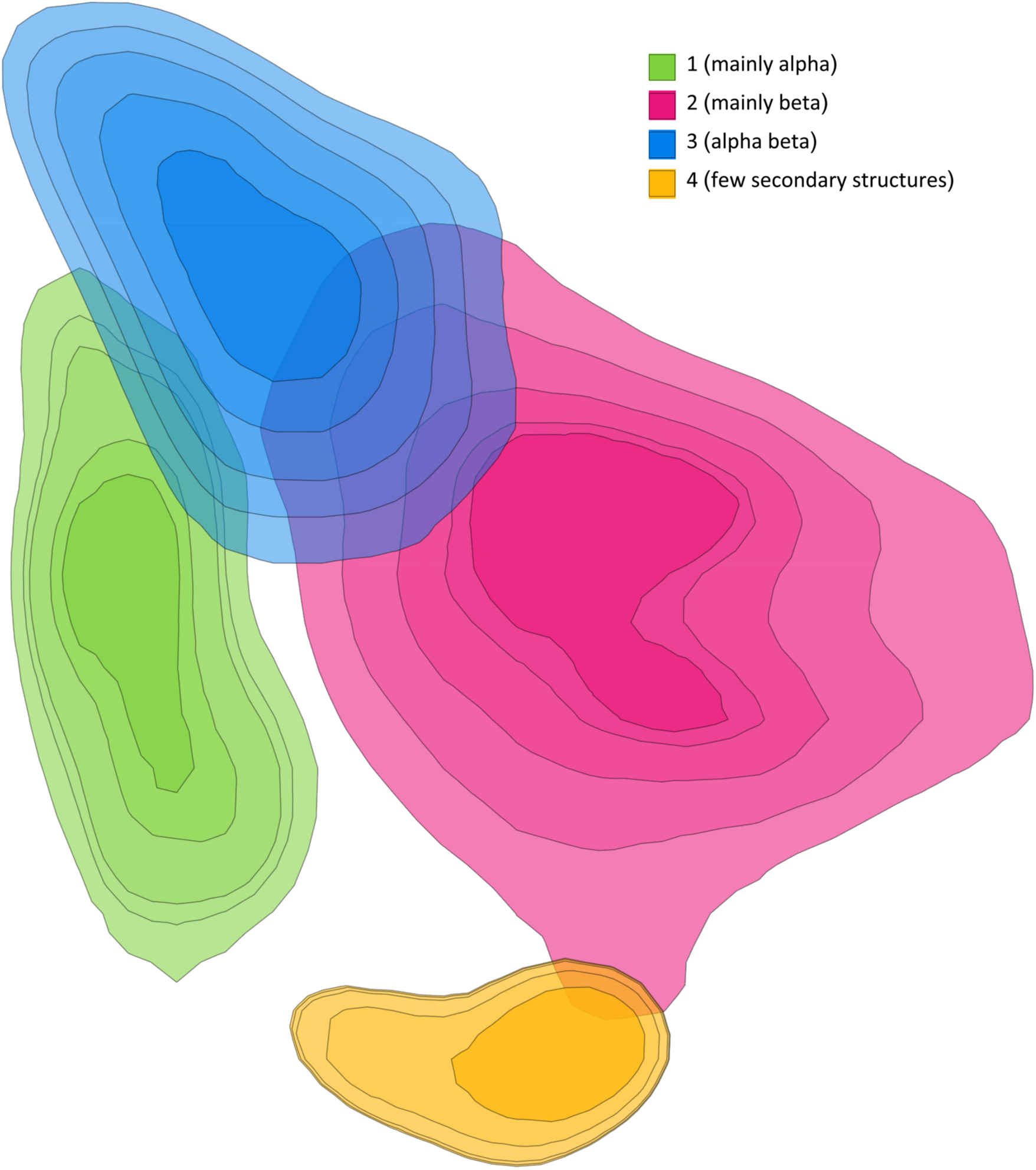
Contour plots of the topmost CATH classes projected onto RGN latent space.

**Fig. S3.**
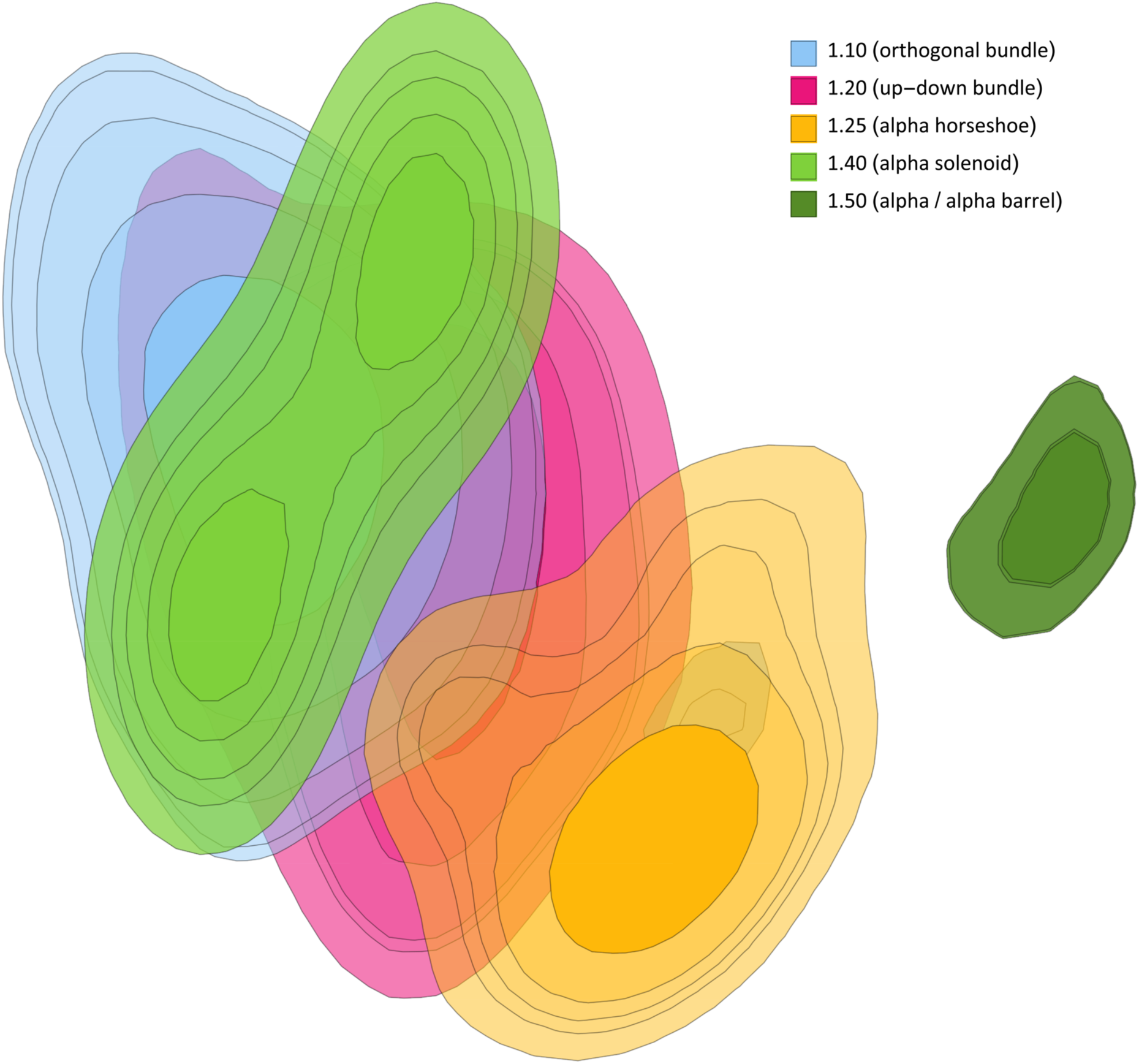
Contour plots of subcategories in the “Mainly Alpha” CATH class projected onto RGN latent space.

**Fig. S4.**
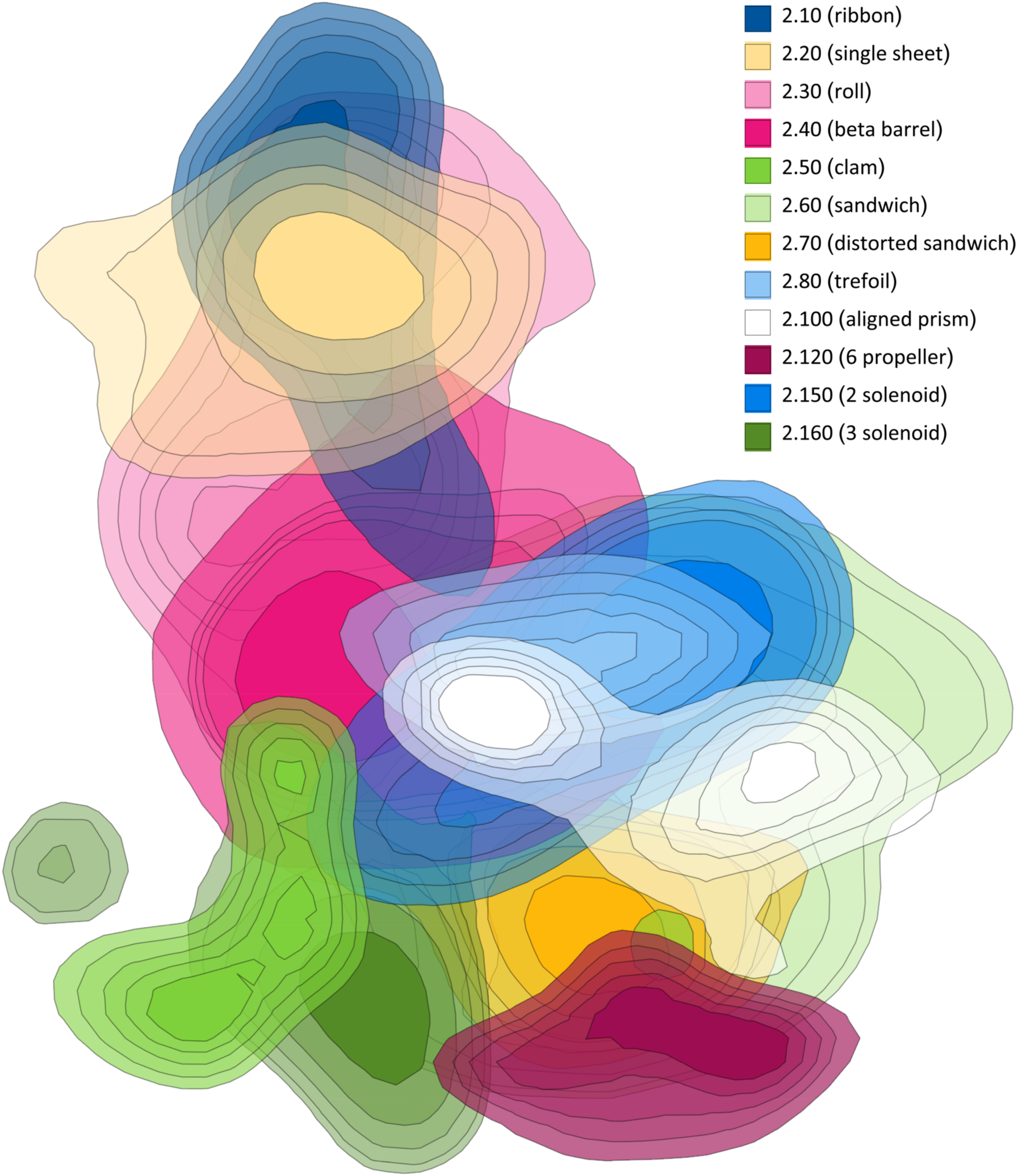
Contour plots of subcategories in the “Mainly Beta” CATH class projected onto RGN latent space.

**Fig. S5.**
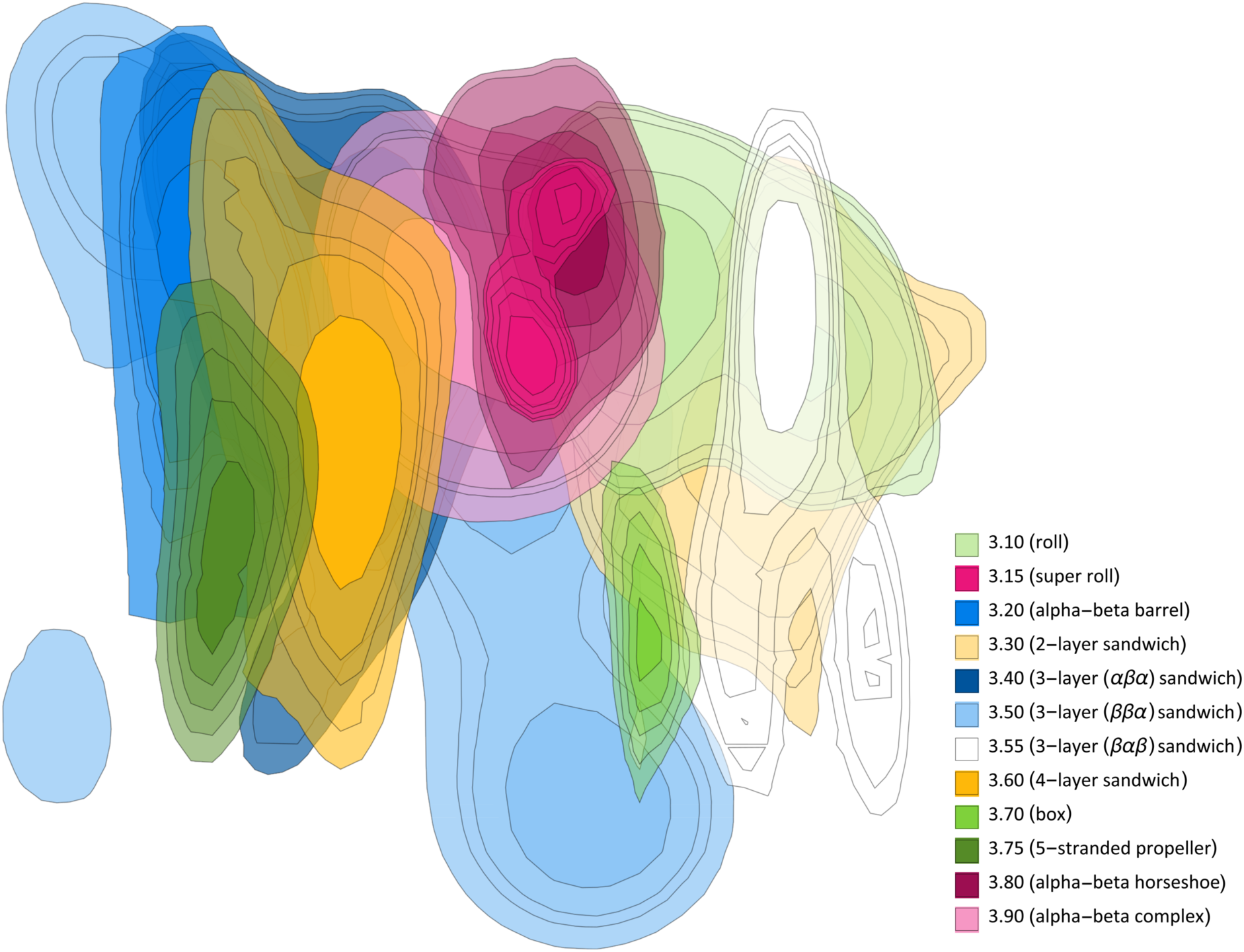
Contour plots of subcategories in the “Alpha Beta” CATH class projected onto RGN latent space.

**Table S1.** Effect of dataset size on RGN accuracy. RGNs trained on ProteinNet (PN) training set X were tested on all CASP test sets subsequent to X (e.g. RGN trained on ProteinNet 7 was tested on CASP 8‐12) to assess the effect of data set size on model accuracy. Numbers shown are differences in average dRMSD (lower is better) relative to RGNs trained and tested on matching data sets (i.e. trained on ProteinNet X and tested on CASP X.)

|  |  | FM (novel folds) test set (Å) |  |  |  |  |  | TBM (known folds) test set (Å) |  |  |  |  |  |
| --- | --- | --- | --- | --- | --- | --- | --- | --- | --- | --- | --- | --- | --- |
|  |  | CASP12 | CASP11 | CASP10 | CASP9 | CASP8 | CASP7 | CASP12 | CASP11 | CASP10 | CASP9 | CASP8 | CASP7 |
| Training set | PN7 | +0.9 | +0.3 | +1.1 | +1.0 | +1.8 | 0 | +1.7 | +1.8 | +0.9 | +1.5 | +0.4 | 0 |
|  | PN8 | +0.6 | +0.2 | +1.2 | +0.3 | 0 |  | +1.4 | +1.0 | +0.2 | +0.9 | 0 |  |
|  | PN9 | 0 | +0.7 | +0.8 | 0 |  |  | +0.6 | +0.6 | 0 | 0 |  |  |
|  | PN10 | +0.5 | +1.2 | 0 |  |  |  | +0.6 | 0 | 0 |  |  |  |
|  | PN11 | +0.2 | 0 |  |  |  |  | +0.1 | 0 |  |  |  |  |
|  | PN12 | 0 |  |  |  |  |  | 0 |  |  |  |  |  |

**Table S2.** Comparative accuracy of RGNs using TM score. The average TM score (higher is better, range is between 0 and 1) achieved by RGNs and the top five servers at each CASP is shown for the novel folds **(left)** and known folds **(right)** categories. Numbers are based on common set of structures predicted by top 5 servers during each CASP. A different RGN was trained for each CASP, using the corresponding ProteinNet training set containing all sequences and structures available prior to the start of that CASP.

|  | FM (novel folds) category (TM score) |  |  |  |  |  | TBM (known folds) category (TM score) |  |  |  |  |  |
| --- | --- | --- | --- | --- | --- | --- | --- | --- | --- | --- | --- | --- |
|  | CASP7 | CASP8 | CASP9 | CASP10 | CASP11 | CASP12 | CASP7 | CASP8 | CASP9 | CASP10 | CASP11 | CASP12 |
| RGN | 0.27 | 0.36 | 0.28 | 0.25 | 0.28 | 0.29 | 0.49 | 0.50 | 0.48 | 0.48 | 0.47 | 0.43 |
| 1<sup>st</sup> Server | **0.33** | **0.37** | **0.32** | **0.30** | **0.29** | **0.35** | **0.72** | **0.72** | **0.71** | **0.69** | **0.66** | **0.70** |
| 2<sup>nd</sup> Server | 0.30 | 0.33 | 0.32 | 0.29 | 0.27 | 0.33 | 0.71 | 0.70 | 0.71 | 0.68 | 0.66 | 0.70 |
| 3<sup>rd</sup> Server | 0.29 | 0.31 | 0.30 | 0.27 | 0.26 | 0.31 | 0.71 | 0.70 | 0.70 | 0.68 | 0.65 | 0.70 |
| 4<sup>th</sup> Server | 0.27 | 0.25 | 0.29 | 0.27 | 0.25 | 0.31 | 0.70 | 0.69 | 0.70 | 0.68 | 0.64 | 0.68 |
| 5<sup>th</sup> Server | 0.24 | 0.24 | 0.28 | 0.26 | 0.22 | 0.30 | 0.68 | 0.69 | 0.70 | 0.67 | 0.64 | 0.68 |

**Table S3.** Validation set milestones for training RGNs. RGN validation performance was monitored during training, and if the shown accuracy milestones were not achieved by the given iteration number, training was terminated and a new model started.

|  |  |  |  |  |  |  |
| --- | --- | --- | --- | --- | --- | --- |
| ProteinNet 7 | Iteration | 1,000 | 5,000 |  |  |  |
|  | dRMSD (Å) | 14 | 13.6 |  |  |  |
| ProteinNet 8 | Iteration | 1,000 | 5,000 | 20,000 | 50,000 |  |
|  | dRMSD (Å) | 13.4 | 13.2 | 12.6 | 12 |  |
| ProteinNet 9 | Iteration | 1,000 | 5,000 | 20,000 | 50,000 | 100,000 |
|  | dRMSD (Å) | 13 | 12.7 | 12.2 | 11.2 | 10.3 |
| ProteinNet 10 | Iteration | 1,000 | 5,000 | 20,000 | 50,000 | 100,000 |
|  | dRMSD (Å) | 12.8 | 12.3 | 11.5 | 10.7 | 9.4 |
| ProteinNet 11 | Iteration | 1,000 | 5,000 | 10,000 | 100,000 | 150,000 |
|  | dRMSD (Å) | 13.7 | 13.5 | 13.2 | 12.1 | 11.4 |
| ProteinNet 12 | Iteration | 1,000 | 5,000 | 20,000 | 50,000 | 100,000 |
|  | dRMSD (Å) | 13.5 | 12.6 | 12.2 | 11.4 | 10.6 |

**Movie S1.**

Backbone trace of an experimental structure of a protein (white) overlaid with RGN‐predicted backbone of the same structure (rainbow colored) as RGN training progresses. Predictions were made every 150 iterations, and the model was trained for approximately 400,000 iterations..

